# Synergy between a shallow root system with a *DRO1* homologue and localized P application improves rice P uptake

**DOI:** 10.1101/2020.11.30.405068

**Authors:** Aung Zaw Oo, Yasuhiro Tsujimoto, Mana Mukai, Tomohiro Nishigaki, Toshiyuki Takai, Yusaku Uga

## Abstract

The development of genotypes and fertilizer management practices that facilitate high phosphorus (P) use efficiency is needed given the depleting phosphorus ore deposits and increasing ecological concerns about its excessive use. Root system architecture (RSA) is important in efficiently capturing immobile P in soils, while agronomically, localized P application near the roots is a potential approach to address this issue. However, the interaction between genetic traits of RSA and localized P application has not been examined. Near-isogenic lines (NILs) and their parent of rice (*qsor1*-NIL, *Dro1*-NIL, and IR64, with shallow, deep, and intermediate root growth angles (RGA), respectively) were grown in flooded pots in a uniform and P-sufficient condition (P_inco_), and with localized P application by dipping seedling roots into P-enriched slurry at transplanting (P-dipping). The P-dipping created an available P hotspot at the soil surface and substantially improved applied P-use efficiency (equivalent biomass at one fifth of application rate of the P_inco_). Further, the *qsor1*-NIL had significantly greater biomass and P uptake than the other genotypes in the P-dipping. The *qsor1*-NIL consistently had a greater root biomass and surface area in the 0–3 cm soil layer, despite that there were no genotype differences in total values and that the other genotypes also reduced their RGAs responding to the P hotspot in the P-dipping. The shallow root system of *qsor1*-NIL facilitated P uptake from the P hotspot. P-use efficiency in crop production can be further increased by combining genetic traits of RSA and localized P application.

## INTRODUCTION

Phosphorous deficiency restricts crop growth, particularly in the tropics, due to the inherently low P content of soils and the high P-fixing capacity of other minerals such as active Al- and Fe-oxides (Walker and Syers 1976). Large amounts of mineral P fertilizer have been continuously applied to overcome low P-use efficiency and achieve high grain yields. Given the finite nature of the P fertilizer resource and increasing ecological concerns about the excess use of P in agricultural systems (Vance et al. 2003; Carpenter and Bennett 2011; Nedelciu et al. 2020), it is vital to investigate sustainable crop production strategies that facilitate the efficient utilization of applied and available P in soils. Such strategies are also critical for the food security of resource-poor farmers with low fertilizer inputs in developing countries (Tsujimoto et al. 2019).

Roots play a pivotal role in exploring immobile P in the soil. An increased root surface area with minimal carbon costs is one strategy, through the formation of finer roots, aerenchyma, and root hairs (Lynch and Ho, 2005; Nestler et al. 2016; Lynch 2019). Changes in root system architecture (RSA) such as the development of surface roots is another root function to adapt to P deficiency, that is so called topsoil foraging, because P is most available in surface soil layers (Lynch and Brown, 2001). This topsoil foraging can be enhanced by a shallower growth angle of axial roots (Lynch and Brown, 2001), adventitious root abundance (Miller et al. 2003), and many/short lateral root branching (Jia et al. 2018). Field-based studies have demonstrated the yield advantages of genotypes with these architectural traits for several crops under P-deficiency (Lynch 2019). Therefore, identification of key root traits and their genetic mechanisms and conferring genes or quantitative trait loci (QTL) should offer avenues for improving P acquisition efficiency in crop breeding (Burridge et al. 2019).

Agronomic approach for improving P-use efficiency includes localized fertilization, which refers to the placement of small amounts of fertilizers nearby the root zone. Several field experiments have demonstrated the positive impacts of localized P fertilization on grain yields and/or fertilizer use efficiencies for crop production (e.g., Vandamme et al. 2018). Our recent study identified that applied P-use efficiency can be substantially improved by dipping seedling roots in P-enriched slurry at transplanting (P-dipping) in severely P-deficient rice fields in Madagascar (Rakotoarisoa et al. 2020). The P-dipping transfers P, with the slurry attached to seedling roots, creating a soluble P hotspot nearby the transplanted roots and facilitating plant P uptake, even under the high P-fixing soils of the tropics (Oo et al. 2020). The use of P-dipping is currently being tested by hundreds of smallholder farmers in Madagascar.

Despite a range of studies in both genetic and agronomic approaches, none have examined how the combination of RSA traits and localized fertilization would affect plant P-use and acquisition efficiencies. In the present study, we aimed to identify the combination effect by using near-isogenic lines (NILs) of *DRO1* and its homologue (*qSOR1*), the major QTLs of rice controlling root growth angle (RGA). The parent variety, IR64, is a high-yielding, modern variety with a relatively shallow RGA with the combination of the nonfunctional allele of *DRO1* and the functional allele of *qSOR1*. The *Dro1*-NIL, developed by Uga et al. (2013), has a relatively deep RGA with the combination of functional alleles of both *DRO1* and *qSOR1*. The *qsor1*-NIL, developed by Kitomi et al. (2020), has a shallower RGA than IR64, with the combination of nonfunctional alleles of both *DRO1* and *qSOR1*. We hypothesize that P-dipping, creating the P hotspot at the soil surface, will have a positive interaction with the shallow root system in rice. By understanding the interaction, further research can be expected to improve applied P use efficiencies by designing RSA traits for localized fertilizer application techniques.

## RESULTS

Localized P application via P-dipping (P_dip_) achieved equivalent biomass and P uptakes at one fifth of the application rate of uniform P incorporation (P_inco_) (Fig. 1). The ANOVA detected consistent and significant interactions between genotype and P treatment for shoot biomass and P uptakes at both 21 days after transplanting (DAT) and 42 DAT. In the P_dip_ treatment, *qsor1*-NIL consistently had greater shoot biomass and P uptake than *Dro1*-NIL. In contrast, in P_inco_, *Dro1*-NIL tended to have greater shoot biomass and significantly greater P uptakes than the other genotypes at 42 DAT. Applied P-use efficiency (calculated as the ratio of shoot P uptake at 42 DAT to the amount of P applied) increased from 3.4% to 16.2% for IR64 by changing the P application methods from P incorporation to P-dipping and further increased to 20.0% by using *qsor1*-NIL (data not shown).

**Fig. 1.**
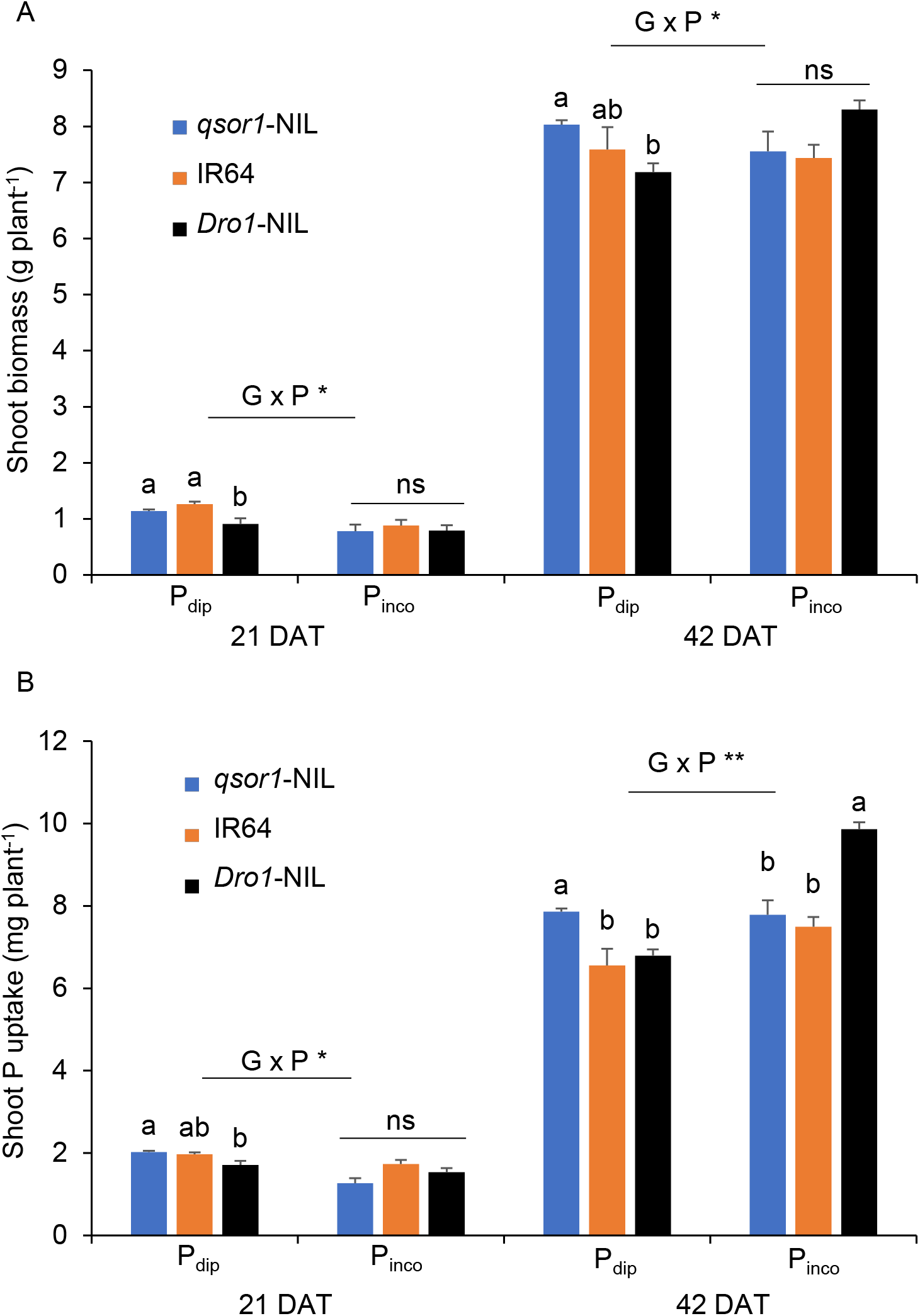
Shoot biomass (A) and shoot P uptake (B) of rice genotypes as affected by different P application methods (P incorporation (P_inco_) of 500 mg P_2_O_5_ box^−1^ vs. P-dipping (P_dip_) of 90 mg P_2_O_5_ box^−1^) at 21 days after transplanting (DAT) and 42 DAT. Different letters and ns within each treatment indicate significant and non-significant differences, respectively, among genotypes at 5% using Tukey's HSD test. Error bars represent the standard error of replications. The * and ** indicate that the interaction between genotype (G) and P application method (P) are significant at P < 5% and P < 1%, respectively.

The RSA traits among genotypes were consistent under P_dip_: the RGA was the shallowest in the order of *qsor1*-NIL > IR64 > *Dro1*-NIL at both 21 DAT and 42 DAT (Fig. 2). As a result of the RGA differences, *qsor1*-NIL developed a large proportion of root biomass and root surface area in the 0–3 cm layer and little in the 14–28 cm layer. In contrast, *Dro1*-NIL distributed a relatively large proportion of root mass in the 14–28 cm layer. For instance, at 21 DAT, *qsor1*-NIL developed 50.3% of the root mass in the 0–3 cm layer and only 2.0% in the 14–28 cm layer while these proportions were 32.7% and 10.3% for *Dro1*-NIL. The root distribution pattern of IR64 was intermediate between *qsor1*-NIL and *Dro1*-NIL. The trend in RSA among genotypes were the same in P_inco_ while IR64 and *Dro1*-NIL tended to have deeper RGAs than those in P_dip_ (Fig. 3). The RGAs of *qsor1*-NIL, IR64, and *Dro1*-NIL at 21 DAT were 7.1°, 23.6°, and 33.3° in P_dip_ and 5.0°, 39.8°, and 52.2° in P_inco_.

**Fig. 2.**
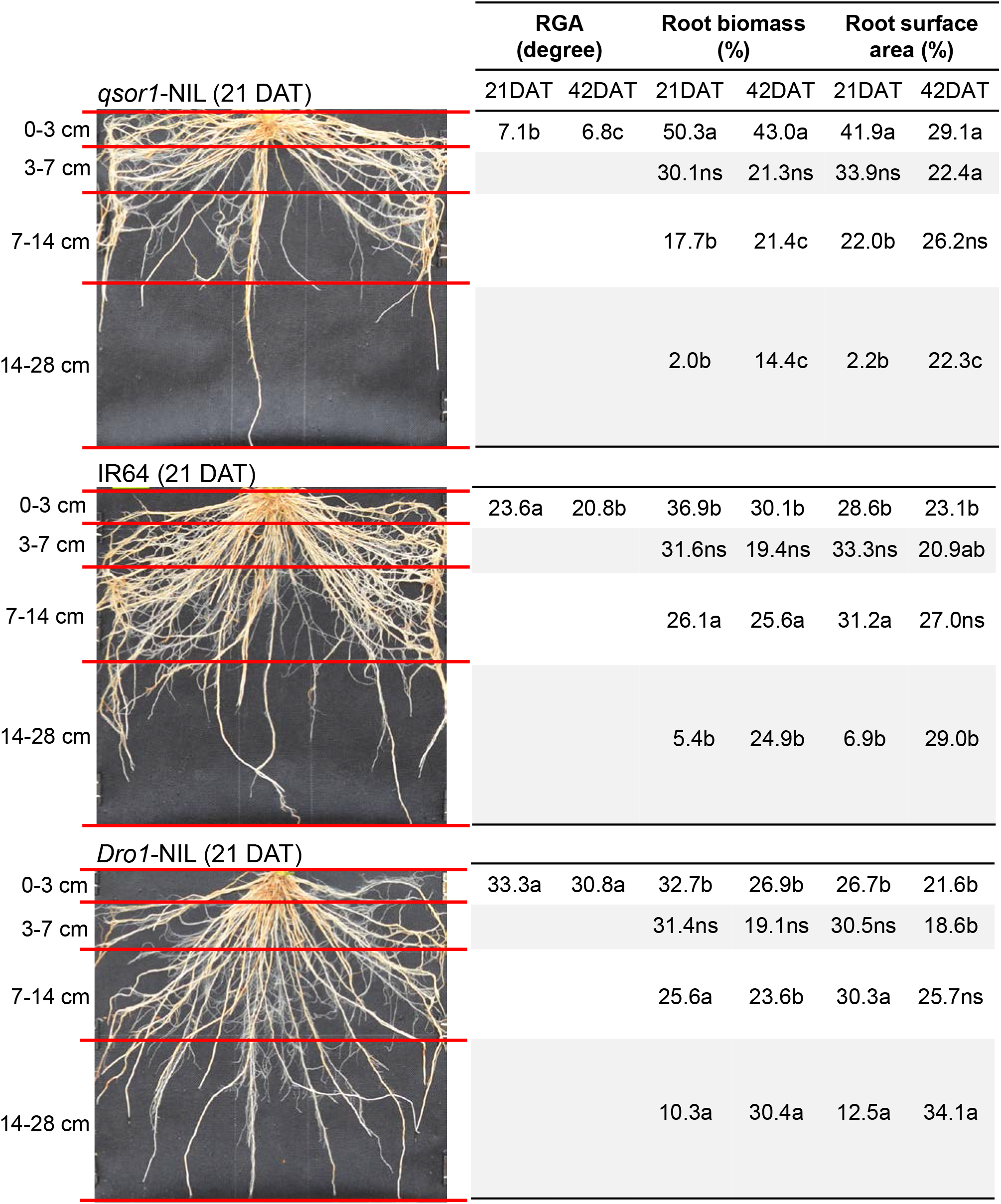
Root growth angle (RGA) and proportions of root biomass and root surface area in different soil layers of *qsor1*-NIL, IR64, and *Dro1*-NIL at 21 days after transplanting (DAT) and 42 DAT under the P_dip_ treatment. Different letters in the same soil layer indicate significant differences among genotypes at 5% of Tukey's HSD test. ns: not significant at 5% level.

**Fig. 3.**
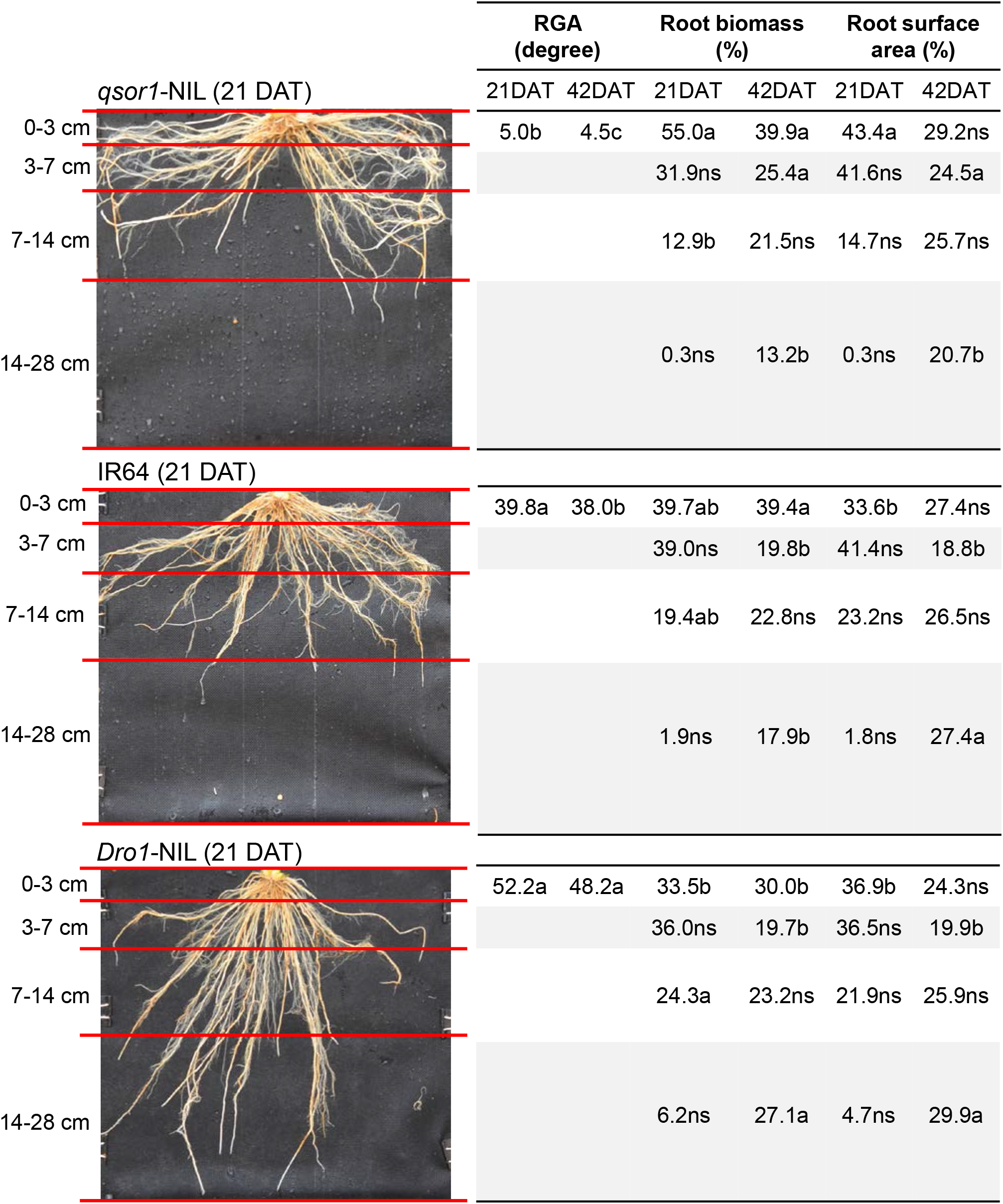
Root growth angle (RGA) and proportions of root biomass and root surface area in different soil layers of *qsor1*-NIL, IR64, and *Dro1*-NIL at 21 days after transplanting (DAT) and 42 DAT under the P_inco_ treatment. Different letters in the same soil layer indicate significant differences among genotypes at 5% of Tukey's HSD test. ns: not significant at 5% level.

By reflecting the differences in RSA, *Dro1*-NIL had a greater root biomass, greater root surface areas, and longer lateral and nodal root length than the other genotypes in the 14–28 cm layer (the difference was only significant vs. *qsor1*-NIL), despite its significantly lower values in the total for these parameters at 21 DAT (Fig. 4). At 42 DAT, there were no significant differences in the total values of these parameters except nodal root length, whereas genotype root distribution patterns were retained within each soil layer: the *qsor1*-NIL had significantly greater root mass, greater root surface area, and longer nodal root length than *Dro1*-NIL in the 0–3cm layer and vice versa in 14–28 cm (Fig. 4). IR64 was intermediate for these parameters in both the 0–3 cm and 14–28 cm layers.

**Fig. 4.**
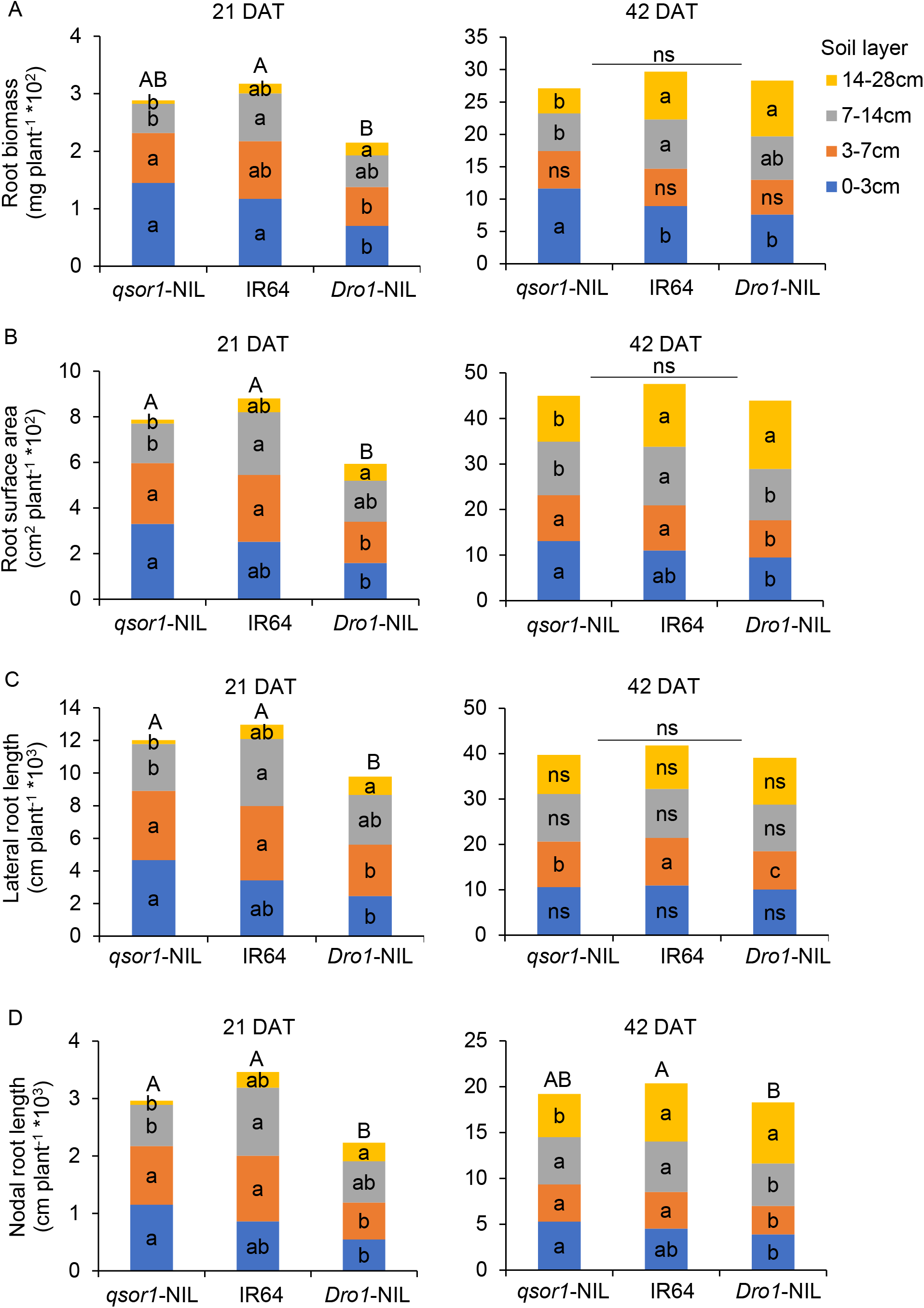
Root development in different soil layers and in total of all layers at 21 days after transplanting (DAT) and 42 DAT under the P_dip_ treatment. Different small letters and capital letters indicate significant differences among genotypes in these parameters within each soil layer and in total of all layers, respectively, at 5% of Tukey's HSD test. ns: not significant at 5% level.

Soluble P concentrations in soils were averaged across genotypes because there were no significant genotype differences in any sampling times or sampling layers. The P_dip_ had a substantially large soluble P concentration at a depth of 3 cm (Fig. 5). The maximum P concentration at a depth of 3 cm for P_dip_ was >100 times greater than the other depths for both P treatments throughout the growing period. In P_dip_, soluble P concentrations were greater at a depth of 7 cm than at 21 cm in the latter growth stages, but apparently the vertical P diffusion from the 3 cm hotspot was relatively small. In contrast, the soluble P concentrations were significantly higher at a depth of 21 cm than at 7 cm in P_inco_ after 28 DAT.

**Fig. 5.**
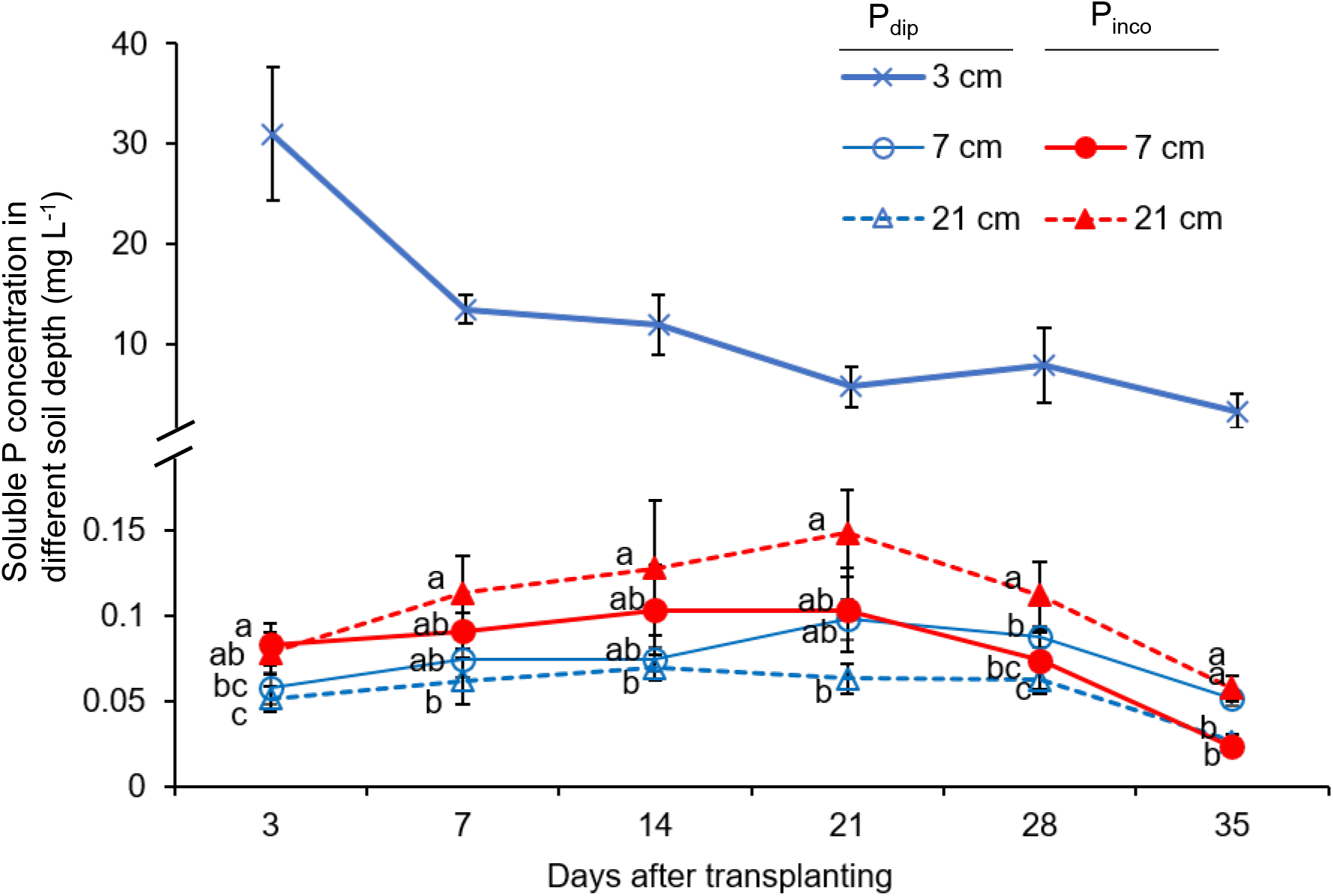
Spatio-temporal variations in soluble P concentration as affected by different P application methods. The cross symbols indicate the value at the 3 cm depth of the P_dip_ treatment. The open and closed circles indicate the value at the 7 cm depth of the P_dip_ treatment and P_inco_ treatment, respectively. The open and closed triangles indicate the value at the 21 cm depth of the P_dip_ treatment and P_inco_ treatment, respectively. Data values are an average of three rice genotypes because no significant genotype difference in soluble P concentration was observed at each sampling time. Error bars indicate standard error of replications. Different letters indicate significant differences at 5% using Tukey's HSD test among different soil depths (7 cm and 21 cm) by P application methods. The observation at 3cm depth was only conducted in the P_dip_ treatment.

## DISCUSSION

The results support the hypothesis that the shallow root system of *qsor1*-NIL has a positive interaction with localized P application via P-dipping and that the combination additively improves applied P-use efficiency for initial rice growth. The other genotypes also reduced the RGA by 16–19° in response to the P hotspot (Fig. 2, 3), yet the synergy with P-dipping was greater in *qsor1*-NIL. This implies that breeding efforts to design the RGA in localized P spots can be more beneficial than relying on the intrinsic root plasticity of each genotype.

Superior P uptake of *qsor1*-NIL with P-dipping is attributable to the greater root biomass and root surface area in the 0–3 cm soil layer where high soluble P is available throughout the growing period. This is most likely the same mechanism as topsoil foraging, prioritizing the root development in the P-rich domains to efficiently capture immobile P in soils. Spatio-temporal P variations in the P-dipping indicate that applied P mobility is highly restricted despite a general understanding that P becomes less immobile under flooded conditions (Turner and Gilliam, 1976), emphasizing the importance of RSA for the localized P acquisition, even under flooded soil culture. The effect of topsoil foraging itself has been reported in several upland crops (Zhu et al. 2005; Miguel et al. 2015; Jia et al. 2018; Sun et al. 2018), but not in rice. Previous studies detected no significant effects of root distribution patterns or RGA for rice P acquisition under P deficiency (e.g., Mori et al. 2016), which may be due to the materials differing not only in root system architecture but in other traits or in more complex screening environments. The present study had an advantage using NILs differing in RGAs otherwise equivalent phenotypes (Kitomi et al. 2020) under non-water-stressed and greatly uneven P availability by P-dipping.

In addition, the present study detected a positive effect of *Dro1*-NIL for P uptake under uniform, P-sufficient conditions. The reason for this positive interaction should be further explored but can be related to consistent P acquisition from the P-rich subsoil layers after the depletion of available P in topsoil layers (Fig. 5). Another potential reason is the more efficient acquisition of other nutrients, such as N, which is vertically more mobile than P. Deep rooting has been reported as a positive trait for N acquisition of upland crops (Lynch, 2019) and also of rice in flooded paddy fields in the latter growth stages (Arai-Sanoh, 2014). In common bean, Rangarajan et al. (2018) postulated that the greater vertical range of roots with deeper RGA and greater number of basal root whorls is advantageous for biomass production when both N and P are deficient. Likewise, dispersed root distribution of *Dro1*-NIL might have benefited from relatively uniform nutrient conditions of the P_inco_ treatment. *Dro1*-NIL had significantly smaller coefficient of variations across soil layers in root biomass at 42 DAT than *qSOR1* (23% vs. 47%), indicating more uniform and dispersed root development.

It should be noted that crop production environments are complex with multiple abiotic stresses, particularly on smallholder farms in developing countries where stress-resilient and nutrient-efficient technologies are most needed. With this respect, field-based experiments to maturity are further required to confirm the combination effect between genetic RSA traits and P fertilizer management practices. The combination of shallow roots and localized P application can never be a silver bullet. A careful selection of field environments where P deficiency is the primary limiting factor is needed to effectively apply this combination, ideally together with the development of bimodal root phenotypes (shallow and deep) against complex growing environments. In rice, *qSOR1* and *DRO1* can be promising genetic resources for the development of such bimodal root phenotypes, as indicated by previous studies (Rose et al. 2013; Uga et al. 2015).

## CONCLUSION

The study provides a significant evidence by using NILs differing in their RGA that a shallow root system has a positive interaction with localized P application nearby the root at transplanting, and the combination substantially improves applied-P use efficiency for initial rice growth. This finding should encourage relevant research focusing not only on physiological root traits or agronomic management approaches, but on their combination to address to the global issue of increasing crop production while minimizing the environmental impacts.

## MATERIALS AND METHODS

### Experimental design and treatments

The experiment was conducted in a greenhouse with an automatic ventilation system at the Japan International Research Center for Agricultural Science (JIRCAS), Tsukuba, Japan. The average daytime and nighttime temperatures during the experiment ranged from 26.2° to 35.8°C and 24.7° to 28.7°C, respectively (Thermo Recorder TR-50U2, T&D Corporation, Japan).

The soil for the experiment was collected from a subsoil layer (40–50 cm in depth) at the JIRCAS Tropical Agricultural Research Front, Okinawa, Japan. The soil was sandy clay and had low pH (H_2_O) of 4.86, low available P content, and high P retention capacity with abundant active Al and Fe oxides. The soil was air-dried and passed through an 8 mm sieve prior to the experiment.

Two different P treatments (sufficient P incorporation (P_inco_) and localized P application via P-dipping (P_dip_)) were factorially combined with three rice genotypes in a randomized complete block design with seven replications. For both treatments, NH_4_NO_3_ and K_2_SO_4_ were mixed with soils and puddled in a bucket at a rate of 220 mg N box^−1^ and 220 mg K_2_O box^−1^ to develop uniform and N- and K-sufficient conditions. For the P_inco_, triple super phosphate (TSP) was added at puddling. Then, the mixed soils were filled into a root box at a rate of 500 mg P_2_O_5_ box^−1^ to develop a uniform and P-sufficient condition. The root box was made of transparent acrylic sheets with a size of 30 cm height × 30 cm length × 3 cm width. The soil was added to the box to a depth of 28 cm.

For the P_dip_ treatment, a P solution was placed in a spot nearby the transplanted root zone to apply the exact amount of P in all boxes. We estimated the amount of P-enriched slurry transferred or attached to seedling roots at transplanting as 90 mg P_2_O_5_ box^−1^ based on our previous study (Oo et al., 2020). After the N- and K-added soil was filled in the root box, 90 mg P_2_O_5_ as TSP dissolved in 20 ml water was injected into the soil at a depth of 3 cm in the center of the root box. On the same day of these P treatments, one 10-day old seedling was transplanted in the middle of each root box and grown under continuously flooded conditions.

### Measurement

Soil solution samplers (DIK-8393, Daiki Rika Kogyo Co. Ltd., Japan) were installed in one side of the acrylic board in the middle of the 3 cm, 7 cm, and 21 cm depths for four out of seven replicates. Soil water samples were collected at 3, 7, 14, 21, 28, and 35 DAT. The samples were analyzed for soluble P concentration using a microplate reader spectrophotometer at an absorbance of 630 nm by following the Malachite Green method (Motomizu et al., 1983).

Three and four replicates were harvested at 21 DAT and 42 DAT, respectively. At each harvest time, shoots were cut at ground level and oven-dried at 70 °C for > 48 h to determine shoot biomass. Shoot P concentration was measured with the molybdate blue method (Motomizu et al., 1983) after dry-ashing at 550 °C for 2 h and digestion with 0.5 M HCl. Shoot P uptake was calculated by multiplying the P concentration and shoot biomass.

After shoots were removed, root samples were collected using pin-board method as per Kano-Nakata et al (2012). In brief, roots were pinned with a 5 mm mesh net and pinboard after which soils were washed off and digital images were taken. The RGA was determined from the digital image as the angle from the soil surface to the shallowest nodal root using ImageJ software (Version 1.52a, NIH, USA). The root system was then divided into 12 compartments or into the center and both sides of the 0–3 cm, 3–7 m, 7–14 cm, and 14–28 cm soil layers to assess spatial root distributions. Root length and surface area of each compartment were measured using Epson Pro-selection X980 Scanner and WinRhizo Pro software (Regent Instruments, Quebec, Canada). Roots were classified as lateral roots (< 0.2 mm) (Sandhu et al. 2016) and nodal roots (0.2 to 2 mm). Roots of > 2 mm were excluded from the analysis, as they were too large for a single root diameter and most likely occurred as a result of a measurement error. After the morphological analysis, root biomass of each compartment was determined by oven-drying at 70 °C for > 48 h.

### Statistical analysis

JMP software (v14.0.0, SAS Institute Inc., Japan) was used to perform the statistical analyses. The treatment means were compared at 5% level of probability using Tukey’s HSD test after the single and/or interaction effects of genotypes and P treatment were confirmed by a generalized linear model.

## Acknowledgements

We thank to Dr. Takuma Ishizaki, Dr. Hiroki Saito, and technical staff members of the Tropical Agricultural Research Front, Ishigaki Island, Japan, for supporting the experimental soil collection. The authors would like to thank Ms. Mayumi Yonemura, Japan International Research Center for Agricultural Sciences, for conducting soil and plant analysis.

